# Accurate localization of linear probe electrodes across multiple brains

**DOI:** 10.1101/2020.02.25.965210

**Authors:** Liu D Liu, Susu Chen, Michael N Economo, Nuo Li, Karel Svoboda

**Affiliations:** Janelia Research Campus, HHMI, Ashburn VA; Baylor College of Medicine, Houston TX; University College London, UK; Boston University, Boston MA

## Abstract

Recently developed silicon probes have large numbers of recording electrodes on long linear shanks. Specifically, Neuropixels probes have 960 recording electrodes distributed over 9.6 mm shanks. Because of their length, Neuropixels probe recordings in rodents naturally span multiple brain areas. Typical studies collate recordings across several recording sessions and animals. Neurons recorded in different sessions and animals have to be aligned to each other and to a standardized brain coordinate system. Here we report a workflow for accurate localization of individual electrodes in standardized coordinates and aligned across individual brains. This workflow relies on imaging brains with fluorescent probe tracks and warping 3-dimensional image stacks to standardized brain atlases. Electrophysiological features are then used to anchor particular electrodes along the reconstructed tracks to specific locations in the brain atlas and therefore to specific brain structures. We performed ground-truth experiments, in which motor cortex outputs are labelled with ChR2 and a fluorescence protein. Recording from brain regions targeted by these outputs reveals better than 100 μm accuracy for electrode localization.

## Introduction

Behavior is produced by organized multi-regional neural circuits (Svoboda and Li, 2018). A major goal of modern neuroscience is to understand behavior in the context of brain-wide maps of neural activity, ideally at the cellular level. Since current methods for recording neural activity in the mammalian brain, such as recordings with extracellular electrodes, sample only a sparse subset of neurons and brain areas in any one experiment, activity maps of the entire multi-regional circuit have to be assembled across recordings from multiple recording sessions and multiple animals performing the same behavior. Precise three-dimensional localization of the recorded neurons in a standardized brain coordinate system is necessary to collate experiments across recording sessions and animals. The somata of recorded neurons are near (< 100 μm) the recording electrodes (Henze et al., 2000); localizing neurons is therefore equivalent to localizing electrodes.

Classical systems neuroscience experiments have often combined neurophysiological measurements with anatomical and and functional mapping to confirm recording locations. For example, studies in the mouse barrel cortex routinely focus on an identified barrel column, which processes information from one corresponding whisker (Petersen, 2019; Yu et al., 2016). Individual barrel columns are recognizable in histological preparations as a ring-like arrangement of cell bodies (the barrel). Small electrolytic lesions can be used to mark the tissue near the electrode for localization in histological material (Sofroniew et al., 2015). In addition, deflection of one whisker excites neurons mainly in the corresponding barrel column, which can be used to identify specific barrels during *in vivo* recordings (Petersen, 2019; Yu et al., 2016). Similar approaches are widely used in recordings from other brain regions that have been deeply explored using anatomical and/or physiological mapping techniques, such as the sensory thalamus and visual cortex (Hubel and Wiesel, 1959).

However, most of the mammalian brain has not been analyzed at comparable levels of detail, and many brain areas do not have finely mapped sensory or motor maps, nor do they contain clear cytoarchitectural features, such as barrels, that could be used for alignment of neurophysiological measurements across multiple brains. Recently developed silicon probes present additional challenges for localization, because they have large numbers of recording electrodes distributed over long linear shanks (Buzsaki, 2004). Specifically, Neuropixels probes (Jun et al., 2017a) have 960 recording electrodes distributed over 9.6 mm shanks. Because of their length, Neuropixels recordings in rodents naturally span multiple brain areas (Allen et al., 2019; Jun et al., 2017a; Steinmetz et al., 2019). Neuropixels probes do not have the electronic circuits to pass the large currents that are required to produce electrolytic lesions. In larger animals it has been possible to localize electrodes in the intact brain using X-ray, MRI, or ultrasound imaging (Cox et al., 2008; Glimcher et al., 2001; Matsui et al., 2007). However, these methods require specialized instruments and have limited resolution and contrast. These methods are also difficult to combine with acute recordings in head-restrained mice, where the probes are inserted and removed from the brain in each experimental session.

Localizing electrodes has been achieved by labeling electrodes with fluorescent dye and post hoc analysis of the recorded tissue using histological methods (DiCarlo et al., 1996; Guo et al., 2017; Jensen and Berg, 2016; Salatino et al., 2017), aided by identifying known electrophysiological features of specific anatomical locations (Allen et al., 2019; Jun et al., 2017a; Steinmetz et al., 2019). However, the accuracy of workflows for localizing electrodes need to be assessed using ground truth experiments.

In addition, recording locations need to be aligned across brains. The structure of each brain differs, even for isogenic animals (Kovacević et al., 2005) and brains deform in an inhomogeneous manner when extracted from the skull and when undergoing various histological procedures. To aggregate recordings across different brains, recording locations have to be precisely localized in individual brains and warped to a standard brain coordinate system.

Here we report a standardized workflow for alignment of Neuropixels probe recordings across individual brains. First, probe tracks are reconstructed in a standardized reference frame. Second, individual electrodes are localized along the track using electrophysiological landmarks. We performed ground truth measurements using optogenetic and fluorescent tagging of axonal pathways. These ground truth measurements reveal the accuracy of electrode alignment to be less than 100 μm.

## Results

### Overview of the workflow

The goal of the pipeline is to localize each electrode (i.e. recording site) on linear probes, and by extension the neurons recorded by that electrode, as accurately as possible in a standardized brain coordinate system (common coordinate framework, CCF v3) (Lein et al., 2007; Oh et al., 2014). The CCF corresponds to a high-resolution image volume of averaged brains based on autofluorescence (Allen Anatomical Template, AAT, http://download.alleninstitute.org/informatics-archive/current-release/mouse_ccf/average_template/) which is used for warping. The AAT is also segmented into brain regions (Allen Reference Atlas, ARA, http://download.alleninstitute.org/informatics-archive/current-release/mouse_ccf/annotation/ccf_2017/) (Dong, c2008.).

Linear probes are studded with a regular pattern of electrodes along the shank (Csicsvari et al., 2003; Jun et al., 2017a). Localization of electrodes requires reconstruction of the probe track using histological methods and mapping the probe locations in the CCF. Furthermore, points along the track, such as the probe tip and a subset of electrodes, are localized along the reconstructed track. This is achieved by identifying the electrodes recording known electrophysiological features that correspond to anatomical landmarks in the Allen Reference Atlas (ARA), which provides brain region annotations for every location in the CCF. The locations of the remaining electrodes are determined by spatial interpolation according to the known interelectrode spacing. The pipeline for localization of electrode locations within the CCF is summarized in Figure 1.

**Figure 1.**
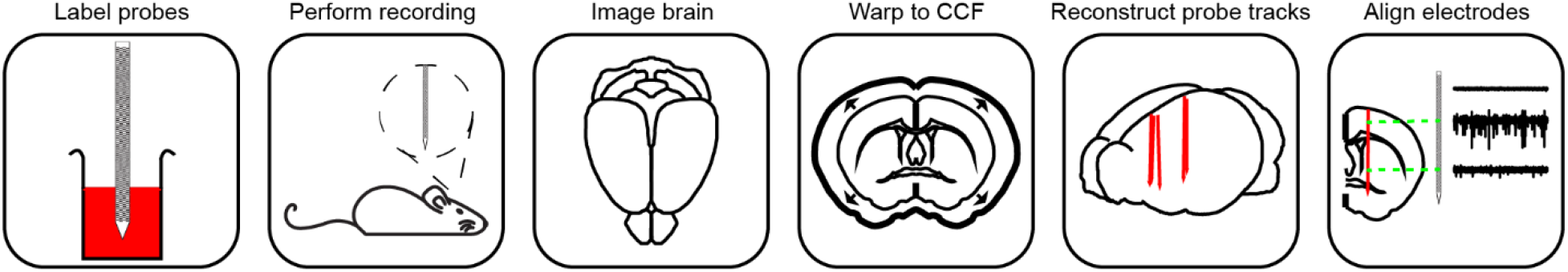
Electrode localization workflow. Before each recording, probes are labelled with a fluorescent dye. After in vivo recordings, the brain is harvested. Fixed brains are cleared and imaged using a light-sheet microscope. The imaged 3D volume is warped to the CCF. The probe tracks are annotated in the 3D volume. Electrodes are localized along the track based on electrophysiological features that correspond to anatomical landmarks.

### Recordings

Acute recordings were made with Neuropixels probes in behaving (Guo et al., 2014a; Inagaki et al., 2018) or untrained awake mice. For the majority of recordings, we used VGAT-ChR2-EYFP mice (Jax 014548) (Zhao et al., 2011), which express ChR2-eYFP in a subset of GABAergic neurons (Figure 2A). Immediately before recordings, probes were coated with the fluorescent dye CM-DiI (Jensen and Berg, 2016). Up to four probes were inserted during each recording session lasting 30-90 minutes. The location of each penetration was recorded with respect to skull landmarks. Penetrations were spaced at least 250 μm apart, which prevented overlap of dye from individual penetrations.

**Figure 2.**
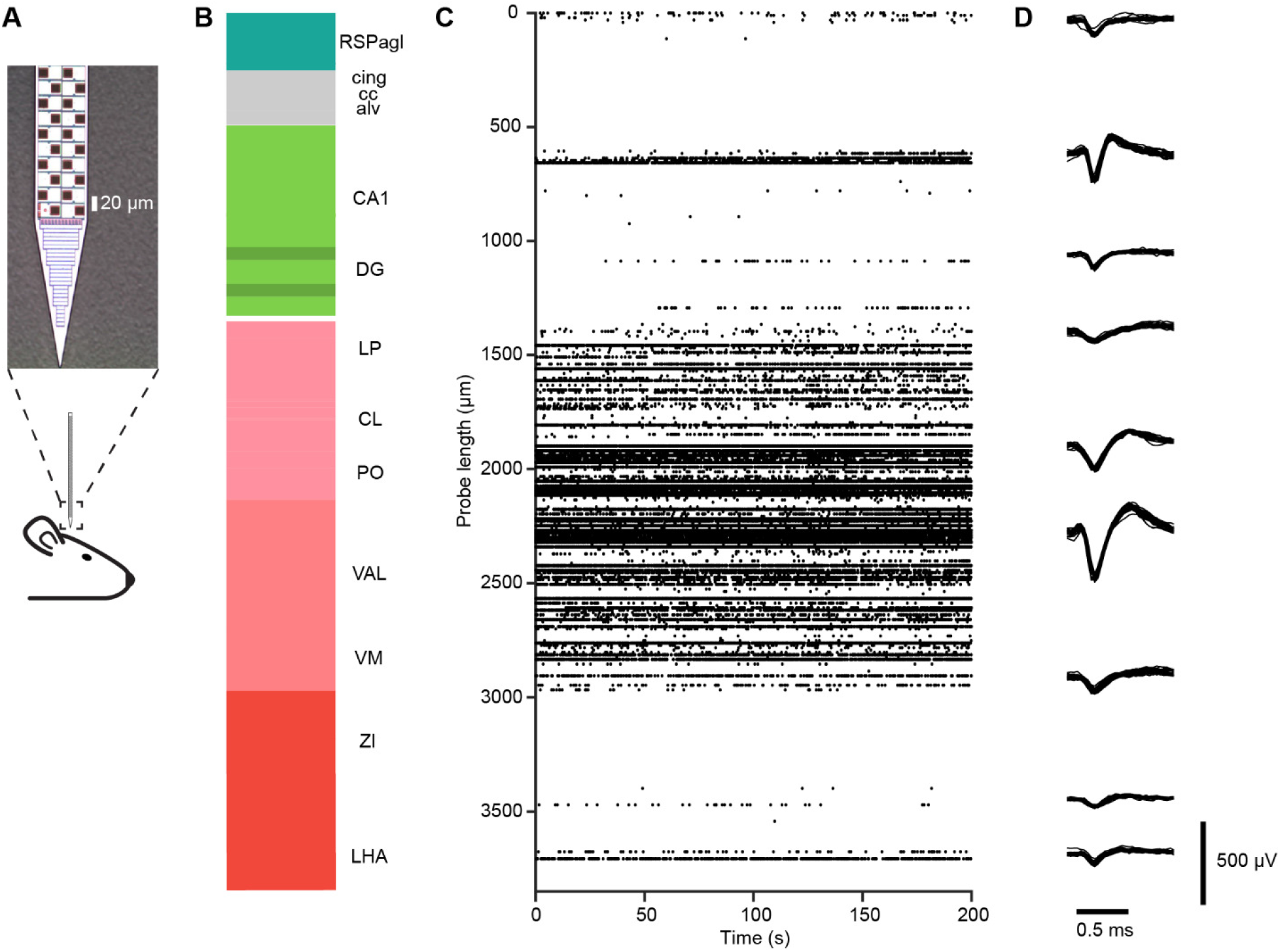
Neuropixels probe recording. (**A**) Schematic of the recording. *Top*, an image of the Neuropixels probe tip showing layout of electrodes. (**B**) An example penetration. The color along the probe track shows Allen Reference Atlas compartments. (**C**) Spike rasters showing multi-unit activity (threshold −70 μV) across electrodes along the probe in (**B**), where 0 is the position of the most superficial electrode. Each event corresponds to a dot on the raster plot. The y-axis indicates the position of the electrodes on the probe. Data from 200 seconds of continuous recording. (**D**) Waveforms from nine example single-units. 25 overlaid waveforms each. The vertical position indicates approximate position along the probe.

Recordings were made simultaneously with 374 electrodes per probe, spanning 3.84 mm of tissue (Jun et al., 2017a). Typically 150-350 units (> 80 μV) were recorded across the brain by each probe (Figure 2B, C). After the last recording session, the animals were perfused, and the brains were carefully extracted and post-fixed (Figure 3A).

**Figure 3.**
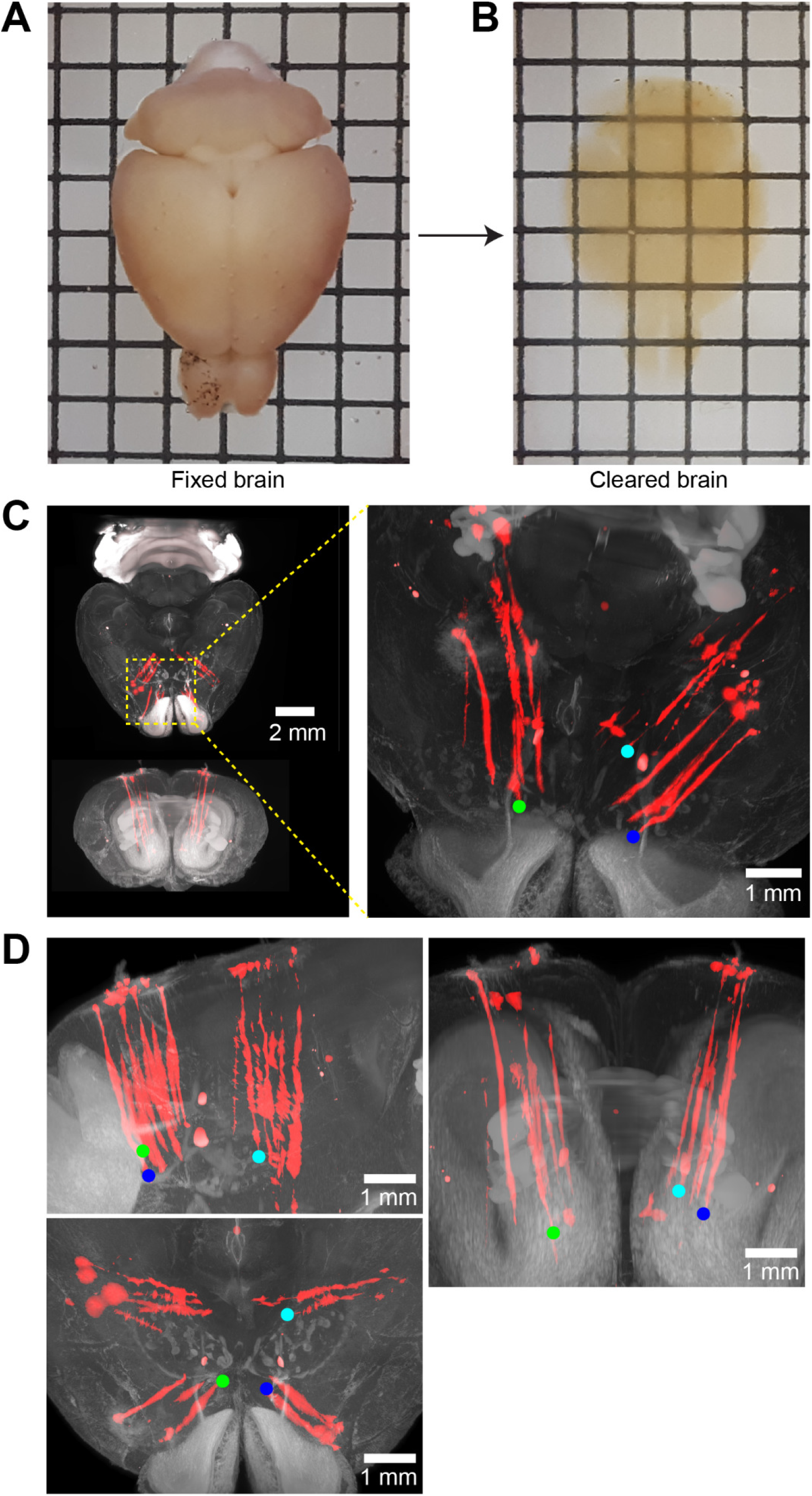
Brain clearing and probe tracks. (**A**) A fixed brain. The spacing between lines in the grid is 2.5 mm. (**B**) Cleared brain in refractive index matched solution. (**C**) Example 3D image volume acquired with light-sheet microscopy. Red shows fluorescence from the CM-DiI labeled probe tracks. Left, horizontal (top) and coronal (bottom) views. Right, zoomed in image of the region around the probe tracks. Image was taken from a different view angle to better show separated probe tracks. (**D**) Images of the probe tracks from different view angles. The tips of three example tracks are marked by colored dots.

### Imaging

We next imaged the fluorescent probe track in the context of brain cytoarchitecture. A variety of imaging methods have been used for whole-brain imaging. Classically the brain is cut into thin (typically 50 μm) sections which are then imaged using standard microscopy or slide scanners. Manual collection and handling of large numbers of sections is labor intensive and error prone. In addition, sections are distorted, complicating assembly of 2D images into coherent 3D volumes that can be aligned to other 3D volumes. These problems can be avoided with microscopes with integrated vibrating microtomes; these microscopes image the tissue in a blockface configuration before cutting, thereby producing high-quality 3D image volumes (Economo et al., 2016; Oh et al., 2014). Indeed, brain volumes imaged in this manner are the basis of the CCF. However, the specialized instrumentation is not widely available and image acquisition across the entire brain is slow.

Instead we used whole-brain clearing methods (Mano et al., 2018) without physical sectioning, in combination with light-sheet microscopy (LSM) (Figure 3B) (Power and Huisken, 2017). We chose a tissue clearing method that results in mechanically robust specimens that are sufficiently transparent for whole brain LSM, preserves the fluorescence of fluorescent proteins, and is compatible with immunohistochemistry (Winnubst et al., 2019) (Methods).

Imaging was done using a commercial LSM with two fluorescence channels (one for the dye; the other for cytoarchitecture). The imaged 3D brain volumes (v3D) show the electrode tracks and distinct cytoarchitecture that can be used for alignment (Figure 3C, D; Supplemental Figure 3.1 {v3Dexp.avi}).

### Three-dimensional templates

We next aligned the v3D to two template brains. First, the AAT is a high-resolution and high-contrast volume of blockface two-photon microscope images, averaged over thousands of brains, where the contrast is based on autofluorescence. The AAT corresponds to a standardized brain coordinate system, the CCF, and an atlas of anatomical structures, the ARA. Second, we constructed a template MRI image volume that better resembles the *in vivo* shape of the mouse brain and allows for more accurate placements of electrode sites.

The appearance of the AAT differs qualitatively from v3D, because of differences in tissue preparation and imaging methods (cf Figure 4A, B). For example, white matter tracts appear dark in the AAT and bright in v3D. In addition, eYFP fluorescence from ChR2-EYFP contributes to the v3D signal (Figure 4A). We therefore used a semi-manual landmark-based method to align brain volumes (Bogovic et al., 2016). Point correspondences between the v3D and AAT were manually determined. To transform the v3D into the AAT space, we used three dimensional thin plate spline interpolation (Duchon, 1977). We first determined the v3D↔CCF transformation by identifying a set of seven landmarks in both the v3D and AAT: the anterior and posterior ends of the corpus callosum along the midline; the meeting point of the anterior commissure along the midline; the genu of the facial cranial nerves to the brainstem in each hemisphere; and indentations from the medial cerebral arteries on the surface of each hemisphere (Table 1; Supplemental Figure 4.1A). We warped individual v3Ds based on this initial set of landmarks landmarks, and then additional landmarks were placed as needed based on visual inspection (Supplemental Figure 4.1B). The warping was performed iteratively after each landmark placement. A 3D volume typically requires 200-300 landmarks to define an accurate transformation (Supplemental Figure 4.1C). A higher density of landmarks was placed around the brain locations containing probe tracks.

**Figure 4.**
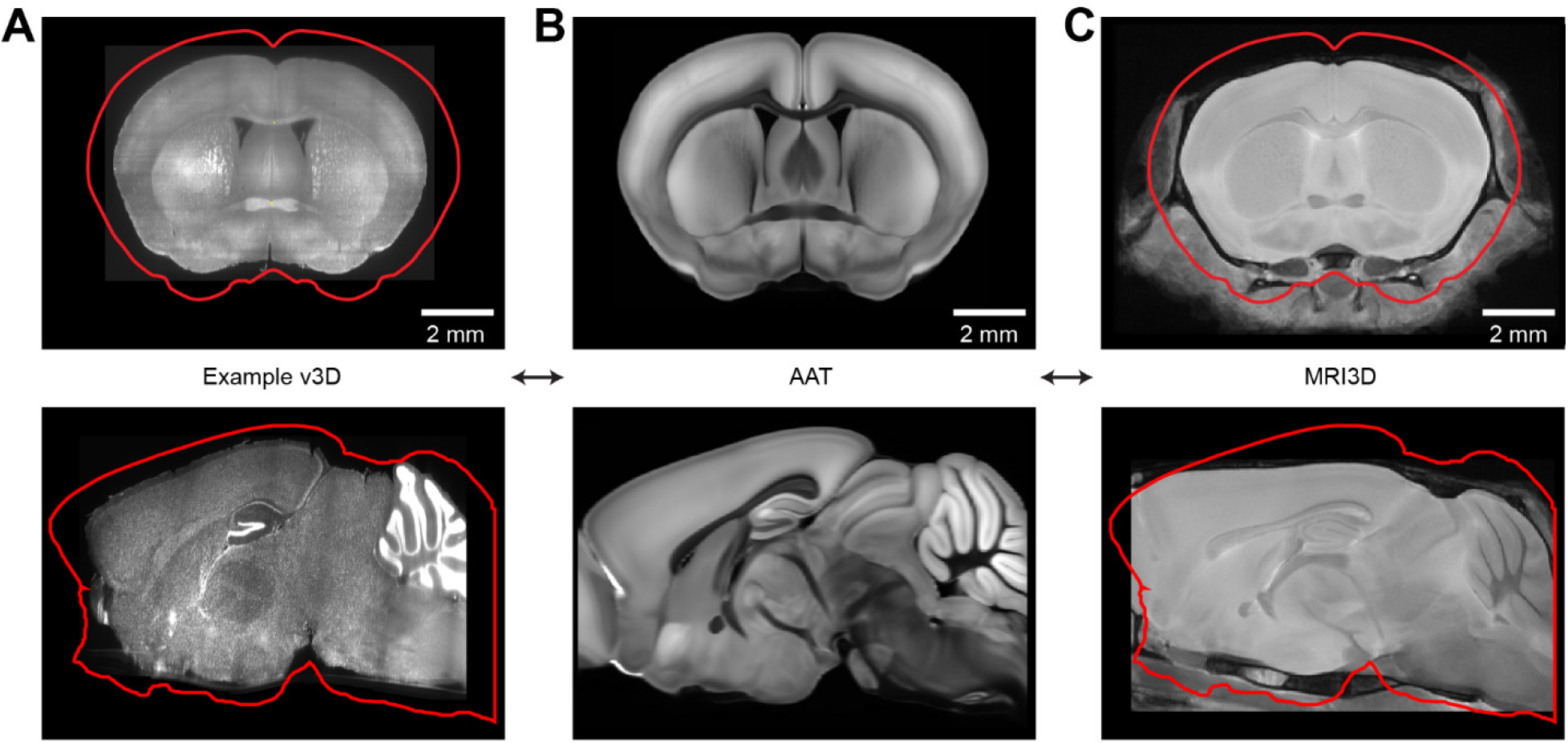
Brain volumes. (**A**) A coronal slice of an example v3D image volume. The red outline corresponds to the Allen Anatomical Template (AAT) Bottom, sagittal slice. (**B**) Same as (**A**), for the AAT. (**C**) Same as (**A**), for the template MRI3D image volume.

**Table 1.** Table of anatomical landmarks in the CCF.

| Landmarks (sFigure 4.1A) | CCF X ($\mu\text{m}$) | CCF Y ($\mu\text{m}$) | CCF Z ($\mu\text{m}$) |
| --- | --- | --- | --- |
| Anterior end of the CC along the midline | 5700 | 3820 | 4240 |
| Posterior end of the CC along the midline | 5700 | 1780 | 7600 |
| The meeting point of the anterior commissure along the midline | 5700 | 5260 | 5160 |
| Indentations for the medial cerebral arteries meeting the hippocampus in DV | 600; 10800 | 4500 | 7720 |
| Genu of the facial cranial nerves to the brainstem | 5100; 6300 | 5100 | 10820 |

The shapes of the 3D image volumes differ across individual mice and differ substantially from the AAT (Figure 4A, B). After multiple recording sessions and penetrations, typically 16 per brain, damage at the insertion sites can cause local deformations. In addition, once extracted from the skull and cleared, fixed brains further deform in a non-uniform manner. For these reasons a relatively high number of landmarks is required.

The AAT was imaged *ex vivo* and is known to be distorted compared to the brain *in vivo*. To warp the v3D into a shape resembling *in vivo* conditions, we imaged VGAT-ChR2-EYFP mice after fixation, but in the skull, using high-resolution MRI (Kovacević et al., 2005; Spencer Noakes et al., 2017). These image volumes are consistent across individual mice (Supplemental Figure 4.2) (Lerch et al., 2008) and are less distorted compared to brains after extraction from the skull (de Guzman et al., 2016; Ma et al., 2008). Individual MRI brains were averaged to obtain the template MRI image volume (‘MRI3D’) (Friedel et al., 2014; Nieman et al., 2018). A comparison of the MRI3D with AAT revealed that the AAT is enlarged compared to the other brain volumes (Figure 4) and distorted in a non-uniform manner (Supplemental Figure 4.3). We established the AAT↔MRI3D mapping using the landmark-based method described above. Thus, any v3D warped into CCF also is automatically aligned to the MRI3D. Because the MRI3D approximately maintains the shape of the brain in the *in vivo* condition, it permits more accurate placement of the electrode sites (below).

### Localization of electrodes

We determined the location of each electrode in the CCF using the following steps. 1.) (Supplemental Figure 1.1: Step 4B) To reconstruct each probe track, we manually placed points at ~200 μm intervals on the centerline of the CM-DiI fluorescence in the v3D (approximately 20 points per penetration). The end of the track was taken to be the point at which DiI fluorescence was no longer visible. A skeleton of the probe track was then determined by linear interpolation between the manually placed points (Figure 5). 2.) (Supplemental Figure 1.1: Step 4C) We projected the probe track into the MRI3D space using the v3D↔CCF and CCF↔MRI3D transformations. Starting at the end of the probe track, we determined the locations of all electrodes along the track using the known inter-electrode spacing. (20 μm for Neuropixels probes). 3.) (Supplemental Figure 1.1: Step 5A,B) We identified characteristic electrophysiological features along the probe that corresponded to known anatomical landmarks (e.g. consecutive electrodes with little activity that correspond to white matter tracts). 4.) (Supplemental Figure 1.1: Step 5C,D) To improve accuracy, we adjusted the electrode positions to align with these electrophysiological features.

**Figure 5.**
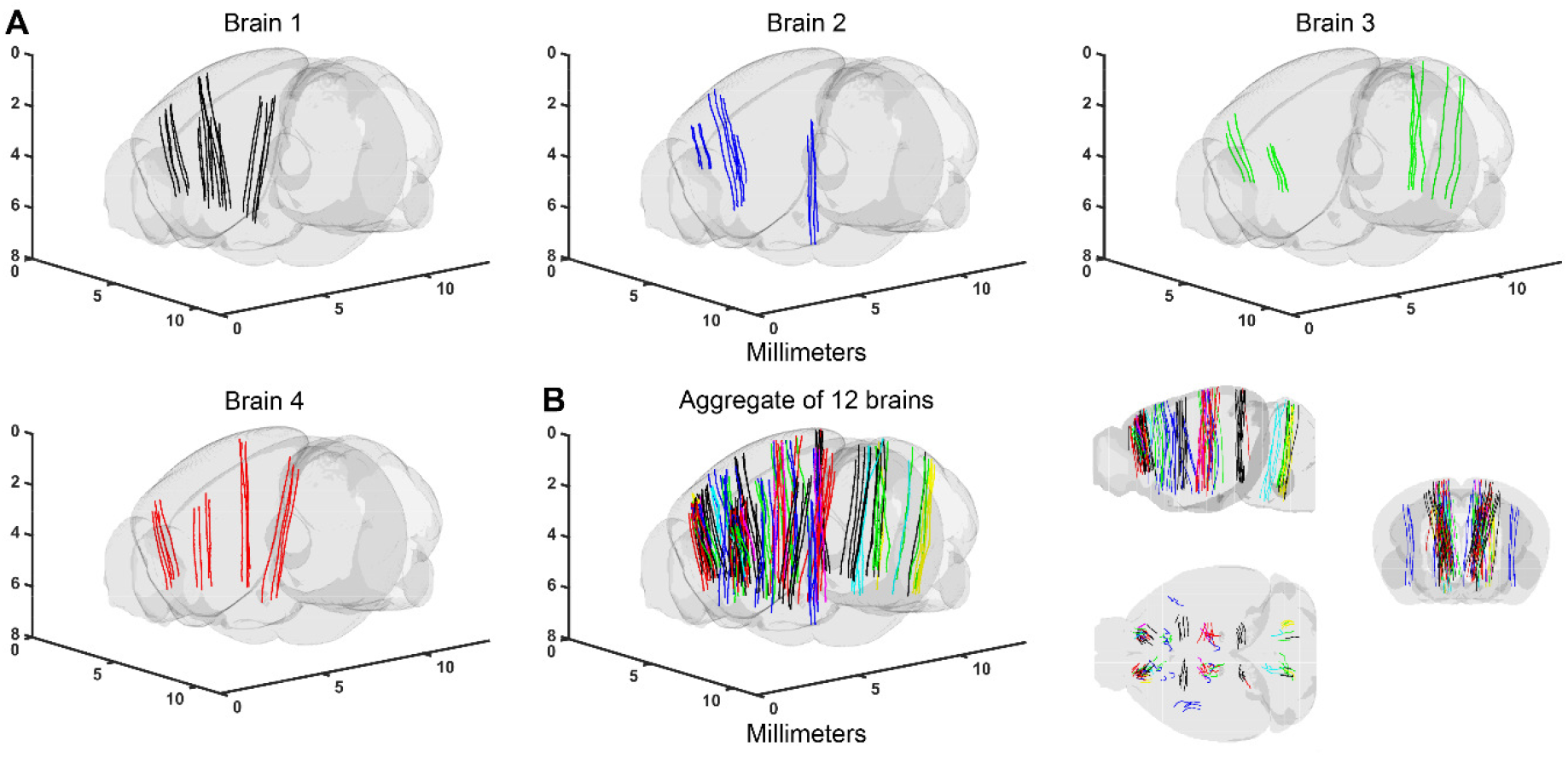
Probe tracks in the CCF. (**A**) Annotated probe tracks in the CCF (four example brains). Brain 1 corresponds to Figure 3C and D. (**B**) Left, aggregate of probe tracks in the CCF. Different colors indicate tracks from different brains). Right, sagittal, horizontal, and coronal views.

Electrode locations were determined within the MRI3D rather than the CCF because the MRI3D is minimally deformed with respect to the *in vivo* brain. We compared the depth of the probe tip in the MRI3D with the depth reading recorded from the micromanipulator. In the CCF the tip location was consistently deeper than the manipulator depth (difference, 1.07 ± 0.37 mm, mean ± SD, 57 penetrations) (at a mean manipulator reading of 3.3 mm), reflecting the fact that the CCF is enlarged compared to the intact brain (Figure 4). In the MRI3D space the mean difference was substantially smaller (0.09 ± 0.26 mm, mean ± SD), reflecting closer resemblance of the MRI3D to the intact brain. The small remaining mean difference between manipulator reading and estimated tip location (0.09 mm, mean) is consistent with dimpling expected at the brain surface after probe insertion (on the order of 100 μm) (O’Connor et al., 2010). The variability in the difference between manipulator position and tip estimate based on histology was substantial across individual penetrations (0.3 mm, SD, MRI3D). This variability reflects uncertainty in the estimate of tip location based on histology: in some experiments the probe tip was brightly labeled with the dye spreading beyond the probe tip, causing an overestimate of probe tip depth. In other experiments the tip was dim, resulting in an underestimate of probe tip depth. This uncertainty makes clear why electrophysiological information is critical to estimate the locations of individual electrodes along the probe track.

After the three-dimensional coordinates of all electrode sites were determined (step 2), we projected these coordinates into the CCF. We then determined the anatomical annotation associated with these coordinates using the Allen Reference Atlas (ARA). We used electrophysiological features recorded on specific electrodes to anchor these electrodes to ARA locations (step 3). Electrophysiological landmarks are anatomical features with recognizable electrophysiological signatures (spiking patterns or local field potentials). Examples include: the surface of the brain, with a sharp transition from low amplitude voltage fluctuations outside of the brain to higher amplitude voltage fluctuations and spikes inside the brain; white matter, such as the corpus callosum (CC), which shows mainly small amplitude axonal spikes compared to the larger spikes in the neighboring gray matter (Figure 6A); ventricles, with no spikes and low amplitude voltage fluctuations (Figure 6A); CA1 layer of the hippocampus, with large amplitude spikes and a phase inversion of the local field potential (LFP) (Buzsáki et al., 2012) (Figure 6B); hippocampus - thalamus border, with low spike rates in the hippocampus, and large amplitude spikes and high spike rates in the thalamus (Figure 6B); vestibular nucleus, showing high spike rates and consistent interspike intervals than in the medulla (Medrea and Cullen, 2013) (Figure 6C).

**Figure 6.**
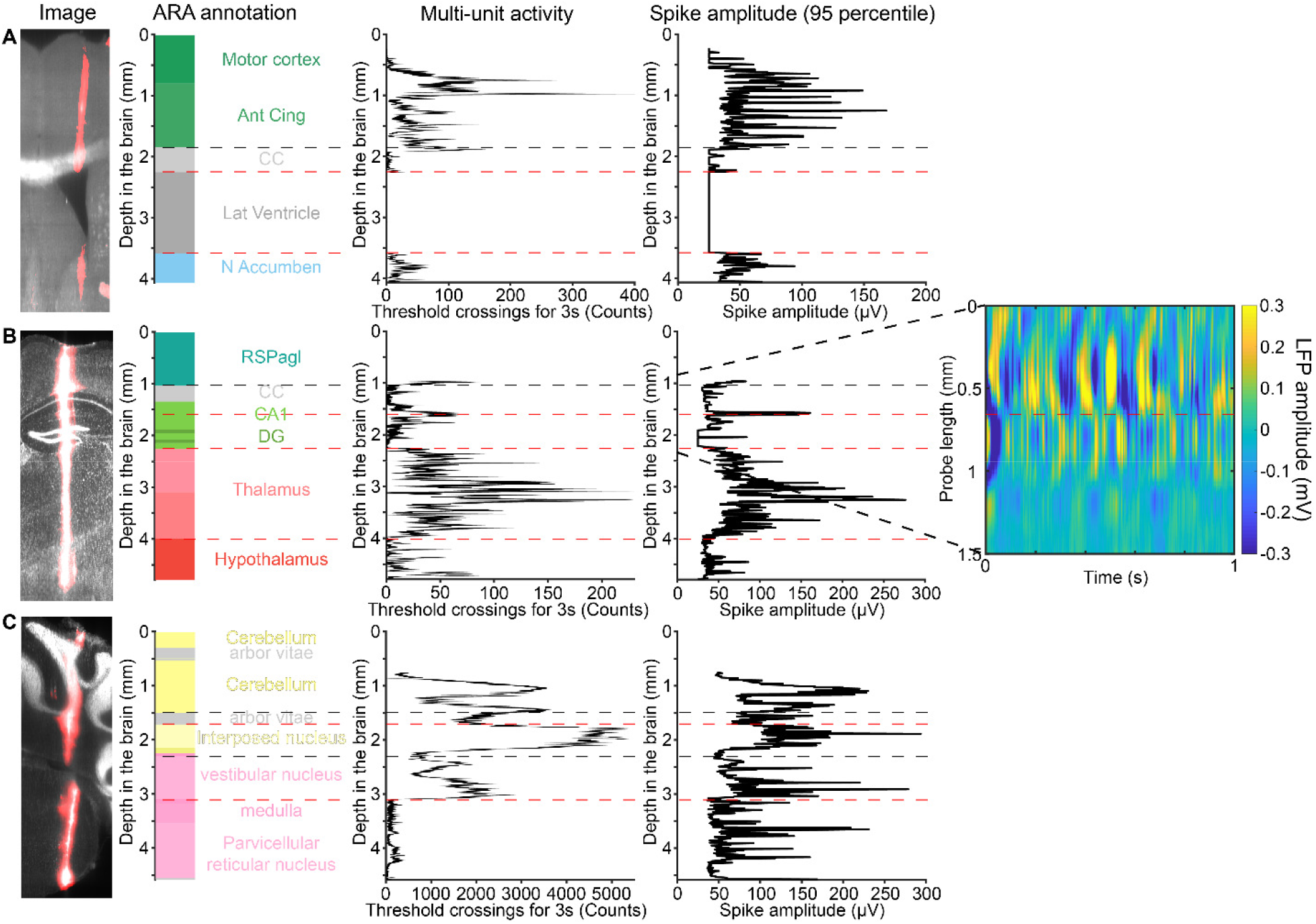
Example electrophysiological landmarks. (**A**) A probe passing through cortex, corpus callosum (CC), and lateral ventricle. The ventricle lacks neural activity. The CC shows small amplitude axonal spikes. (**B**) A probe passes through the CA1 pyramidal cell layer, which shows up as a narrow band of large-amplitude spikes and a phase inversion of the LFP (right insert, raw LFP amplitude). In addition, the borders of the thalamus are marked by the presence of large amplitude spikes. (**C**) A probe passes through the deep cerebellar nuclei (DCN) and the medulla. The transition from the DCN to medulla is marked by a dip in spike rate. The upper portion of the medulla corresponds to the vestibular nucleus, which has high spike rates. The arbor vitae (cerebellar white matter), which mostly lacks multi-unit activity, is not used as an electrophysiological landmark in the electrode localization process, but shows good agreement with the ARA annotation.

The electrophysiological landmarks were then used to localize a subset of electrode sites to ARA compartments or transitions between compartments (Red dashed lines, Figure 6) (step 4). Two or more electrodes need to be localized per probe. The inter-electrode distances in between two electrophysiological landmarks are scaled linearly using a common scaling factor in the MRI3D (because the CCF is enlarged and distorted). The inter-electrode distance above the first electrophysiological landmark or the inter-electrode distance below the last electrophysiological landmark are extrapolated using the nearest scaling factor. The computed electrode tip locations corresponded accurately to electrode depths derived from manipulator readings (difference, 0.08 ± 0.11 mm, mean ± SD).

### Groundtruth experiment

We next performed an independent set of experiments to quantify the accuracy of our procedure for electrode localization. We performed experiments in wild-type C57BL/6J mice. Neurons in the left anterior lateral motor cortex (2.5 mm anterior of bregma, −1.5 mm lateral) were transduced with AAV virus expressing ChR2-eYFP (Guo et al., 2014b; Li et al., 2015). We recorded from downstream brain regions that contained small axonal projections expressing ChR2-eYFP, including a subset of the locations we used as electrophysiological landmarks (based on neural activity outside of photostimulation). Photostimulation of these axons produces phasic neural activity with short latencies on electrodes near the ChR2-eYFP expressing axons. We used this activity to confirm that we have correctly identified key electrophysiological landmarks, such as the white matter. More importantly, the intersection of the eYFP signal in 3D volumes and the probe track provides an independent confirmation of the location of the electrophysiological landmarks.

ALM axons cross the corpus callosum into homotypic ALM in the right hemisphere (Li et al., 2015). Small multi-units events reflecting axonal spikes were visible in extracellular recording (Figure 7B). In response to short optogenetic stimuli (duration 2 ms) we detected a phasic increase in multiunit activity with short latency (2 ms) (Figure 7C).

**Figure 7.**
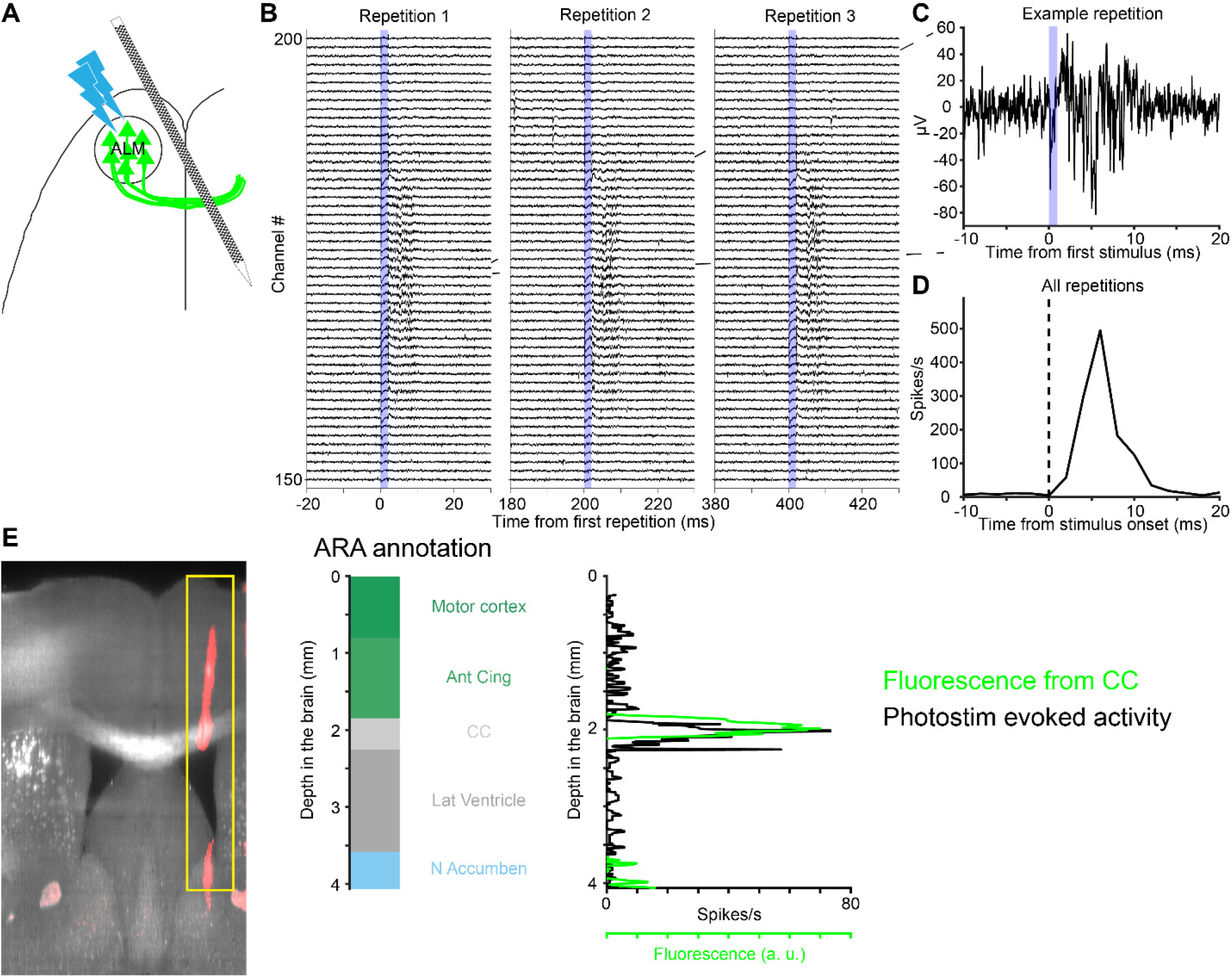
Groundtruth experiment with photostimulating axons in the corpus callosum. (**A**) Schematic of the experiment. ChR2-eYFP was expressed in ALM neurons. A photostimulus was applied over ALM to elicit spikes, while recording from contralateral ALM axons. (**B**) Example voltage traces show evoked multi-unit activity in the CC during three successive photostimuli. Voltage traces were from 50 electrodes around the CC. The blue shading indicates the 2 ms stimulus pulse duration. (**C**) Voltage trace from one electrode in response to one photostimulus. (**D**) Peri-stimulus time histogram of multi-unit events (averaged over 300 repetitions for the channel in C). (**E**) Left, a coronal section showing fluorescence from ChR2-eYFP (light grey) and CM-DiI (red) labeled probe track. Middle, ARA annotation along the probe. Right, intensity of ChR2-eYFP fluorescence (green) and evoked activity (black) along the localized electrode in the CCF. Importantly, the corpus callosum was not used as an electrophysiological landmark in the electrode localization process.

ChR2-eYFP expressing axons also project to the mediodorsal and ventromedial nucleus of the thalamus (MD and VM) (Guo et al., 2017). Photostimulation of ALM evoke postsynaptic responses in the MD and VM neurons (Figure 8B). The responses in the MD and VM are of longer latency (5 ms) (Figure 8C), consistent with the length of synaptic delay.

**Figure 8.**
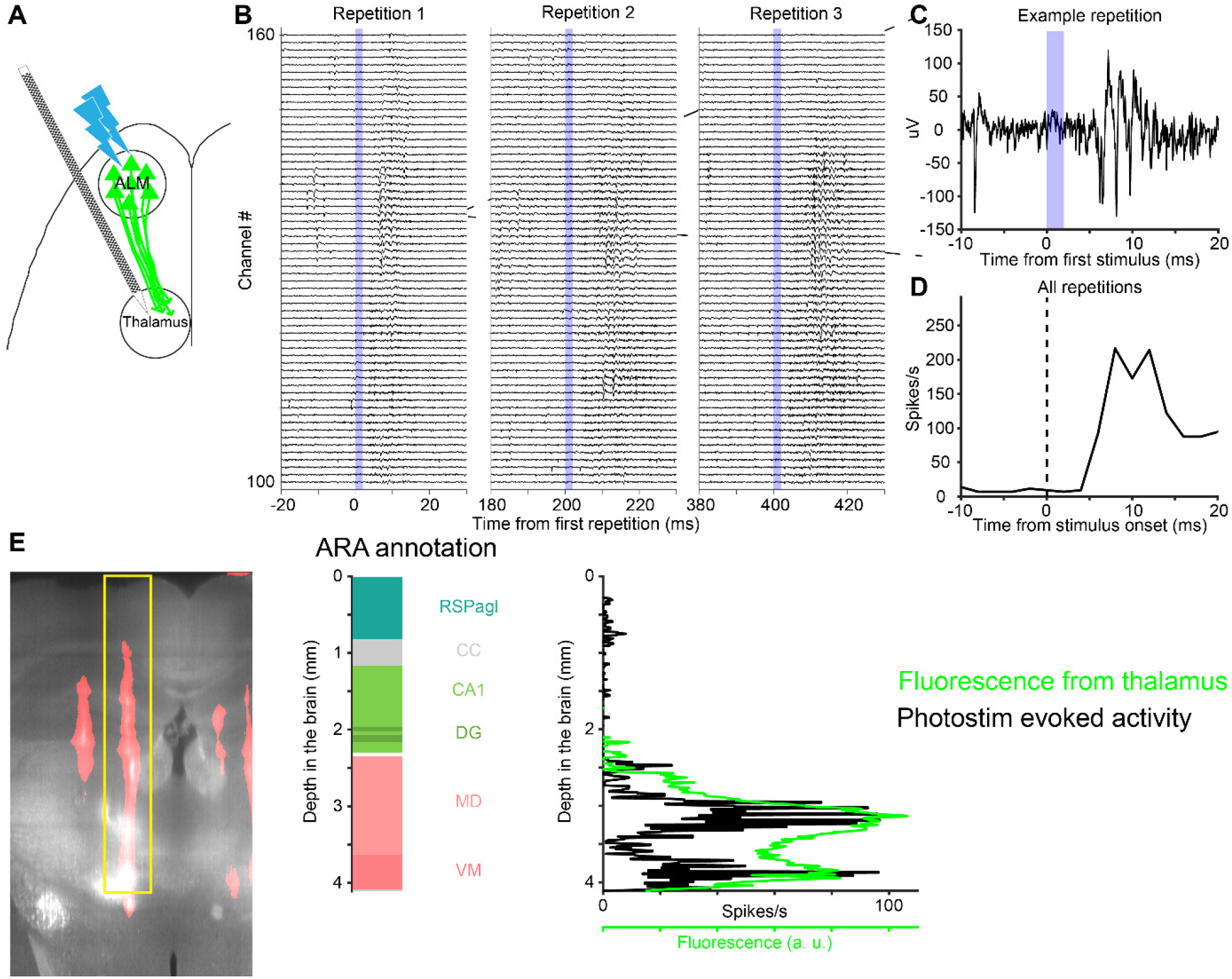
Groundtruth experiment photostimulating axons in the thalamus. (**A**) Schematic of the experiment. Activity was recorded in the thalamus. (**B**) Example voltage traces show multi-unit activity in the thalamus. Same as Figure 7B, but for 60 electrodes in the thalamus. (**C**) Voltage trace from one electrode in response to one photostimulus. (**D**) PSTH of multi-unit events. Averaged over 300 photostimulus repetitions for the channel in C. (**E**) Left, a coronal section showing fluorescence from ChR2-eYFP (light grey) and CM-DiI (red) labeled probe track. Middle, ARA annotation along the probe. Right, intensity of ChR2-eYFP fluorescence (green) and evoked activity (black) along the probe in the CCF. Importantly, the medial dorsal and ventral medial nuclei of the thalamus were not used as electrophysiological landmarks for electrode localization.

The evoked activity confirmed the electrode placement using electrophysiological landmarks. We localized each electrode in v3D and CCF, in which the eYFP fluorescence indicates the ChR2 expression. The profile of the evoked activity on the electrodes resembled the profile of the eYFP fluorescence (Figure 7E and 8E). Quantifying the peak locations of the fluorescence and evoked activity gave us a quantitative description of the accuracy of the alignment procedure (70 ± 56 μm, mean ± SD) (Figure 9, Methods)

**Figure 9.**
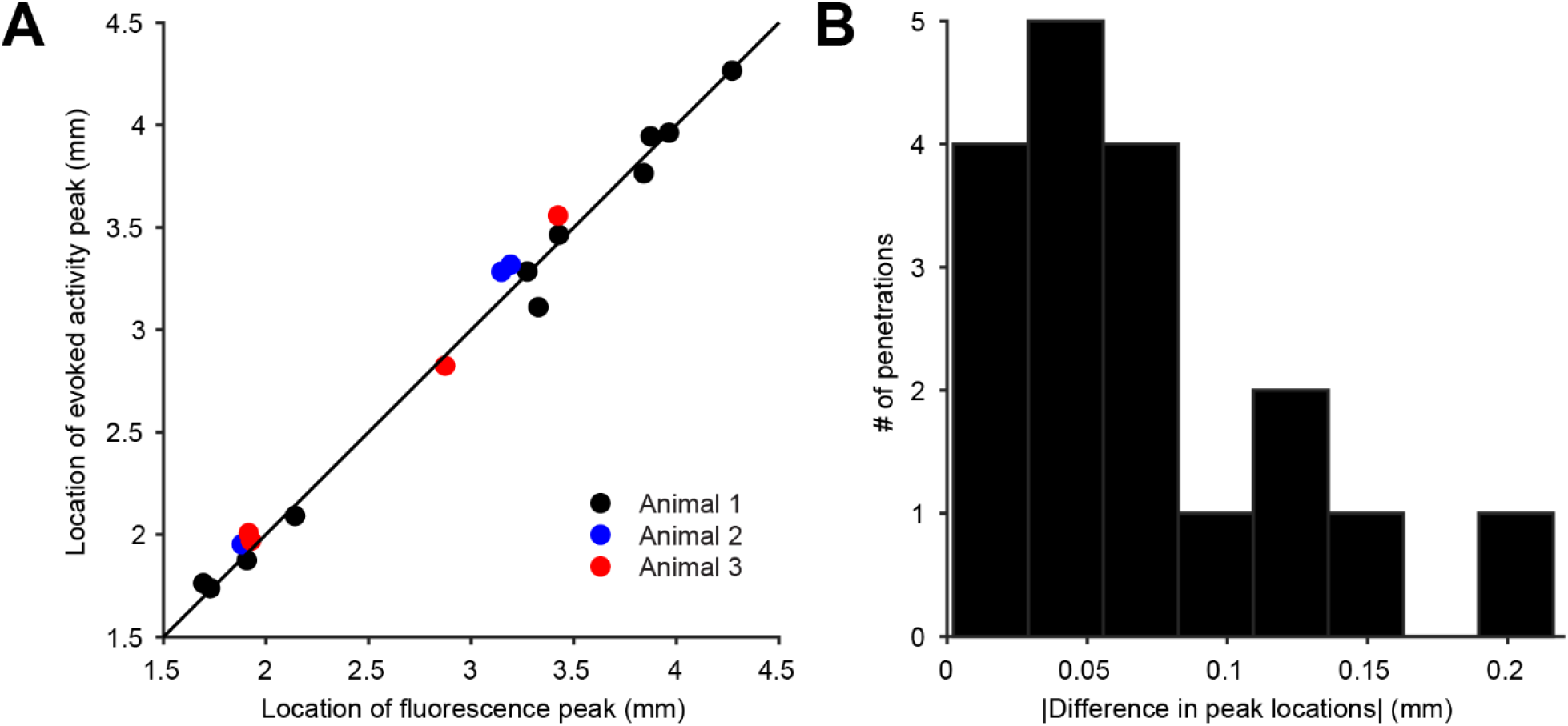
Accuracy of electrode localization as assessed by the groundtruth experiment. (**A**) The peak locations of evoked activity and eYFP fluorescence were estimated using a Gaussian fit (three mice, 15 penetrations). (**B**) The distance between the peak locations of evoked activity and eYFP fluorescence. Absolute value of the difference in peak locations was used to quantify electrode localization accuracy (70 ± 56 μm, mean ± SD).

Using ground truth experiments we evaluated the impact of using the MRI3D for accurate electrode placement. We directly placed electrodes in CCF (without MRI3D), using linear interpolation between electrophysiological landmarks. Direct placement of electrodes in CCF was less accurate compared to placements in the MRI3D (125 ± 63 μm, mean ± SD; *P* = 0.003, two-tailed paired t-test).

## Discussion

We describe a pipeline to localize all electrodes along a linear probe in a standardized mouse brain coordinate system. During recordings, probe tracks were marked with fluorescent dye that persisted in the tissue across multiple experiments spanning weeks. After the experiments, the brain volume and probe tracks were imaged in the intact brain *ex vivo* using a combination of tissue clearing and light-sheet microscopy. The imaged volume was computationally warped to the CCF. Individual electrodes were localized along the probe track based on electrophysiological signatures and other electrodes were assigned by interpolation and extrapolation. Groundtruth experiments indicate that our pipeline has an mean accuracy of 70 μm for localizing electrodes in the CCF.

Neuropixels probes and other large linear probes (Fiáth et al., 2018; Raducanu et al., 2016) sample activity across multiple brain regions. Interpretation of neural activity in terms of neural circuits relies on accurate localization of individual electrodes and thereby the recorded neurons. Several workflows have been described (Allen et al., 2019; Siegle et al., 2019; Steinmetz et al., 2019), but groundtruth experiments assessing the accuracy of electrode localization are lacking. We took advantage of the ChR2-EYFP in bundles of axons and compared the locations of ChR2-EYFP expression as judged by fluorescence with light-evoked activity measured with electrodes that were registered to the CCF. This method can be readily applied to assess the accuracy of other electrode localization pipelines using different types of linear probes, different 3D imaging methods, and possibly in different species.

The CCF is distorted compared to the intact brain: the brain is enlarged and locally sheared, especially around the ventricles. We obtained an MRI3D image stack for our laboratory mice. Mice were matched by strain, age, sex and experimental condition (water restriction). Furthermore, the MRI3D was imaged inside the skull which prevented enlargement and local distortion of the brain. The MRI3D resembled the *in vivo* conditions of the recorded brains (de Guzman et al., 2016). The placement of electrodes in the MRI3D was more similar to the electrode spacing in the intact brain, thereby, leading to more accurate estimate of electrode locations. Groundtruth experiments show that inclusion of the MRI3D improved the accuracy of electrode localization (accuracy from mean 120 μm to 70 μm).

Brain shapes differ across different mouse strains and also depend on the sex of the animals. For example, we note that the brains of our laboratory mice were approximately 10% smaller than C57Bl/6J mice previously imaged under the same conditions using MRI (Dorr et al., 2008; Steadman et al., 2014). Using the workflow introduced here may require acquisition of additional MRI volumes for different mouse strains.

Histological methods by themselves were not sufficient to localize electrodes (errors up to 0.3 mm, SD). Electrophysiological signatures are necessary for anchoring sites along the probe, followed by interpolation and extrapolation outside of the electrophysiological landmarks (Peters et al., 2019; Siegle et al., 2019; Steinmetz et al., 2019). Our work further extends this concept to multiple brain regions and improves the procedure by placing the electrodes in the MRI3D space.

One potential limitation when applying the electrode localization pipeline at large scale is the manual landmark placement for warping individual v3Ds to the CCF. The difference in fluorescence between the v3D and the CCF prevents standard fully automated alignment methods (Kuan et al., 2015). This issue could be overcome by constructing an in-house template for the v3D brains using the same tissue preparation and imaging conditions. The template can be carefully warped to the CCF using the manual procedures described. Each individual v3D can then be warped to the template automatically instead of warping to the CCF.

Another manual step is the selection of the electrophysiological landmarks for interpolation of electrode sites. The electrophysiological landmarks identified here tend to be robust in the regions of interest (Small SD in the Multi-unit activity compared to the mean fluctuation in Figure 6). Similar probe tracks traversing through similar brain regions have similar electrophysiological landmarks. Currently, the procedure of localizing electrophysiological landmarks requires manual selection of the electrophysiological features by the experimenter (Table 2). This step requires knowledge of the underlying anatomy. Detailed analysis of electrical activity in different brain regions might allow automated procedures. In the future, automating 1) warping of v3D to template brains, 2) reconstruction of probe tracks, and 3) anchoring of electrodes using defined electrophysiological landmarks will greatly accelerate the electrode localization pipeline.

**Table 2.** Table of electrophysiological landmarks.

| Landmarks (Figure 6) | Electrophysiological feature |
| --- | --- |
| Ventricle/brain surface | Lack of spiking activity |
| Fibre tract | Reduced activity and smaller spike amplitude compared to cortical regions |
| CA1 pyramidal cell layer | Higher activity and larger amplitude spikes than other parts of the hippocampus; LFP phase inversion |
| Hippocampus to thalamus boundary | Higher activity and spike amplitude in thalamic nuclei than hippocampus |
| Vestibular nucleus | Higher activity than other parts of the medulla |
| Medial superior olive | Sound related-activity in auditory tasks |

## Methods

### Surgeries and Animals

All animal experiments adhered to the guidelines set by the Janelia Research Campus Institutional Animal Care and Use Committee. Nine VGAT-ChR2-eYFP (JAX 014548, >P60, all male) and three wildtype C57BL/6 mice (>P60, 1 male and 2 female) were used in this study. The details of the surgery procedure can be found elsewhere (Guo et al., 2014b) (dx.doi.org/10.17504/protocols.io.bcrsiv6e). Briefly, mice underwent stereotaxic surgery to implant headbars for head-fixation for electrophysiological recordings. The skull was made clear for photostimulation experiments (Guo et al., 2014b). The skin and periosteum were removed, and a thin layer of cyanoacrylate (Krazy glue) was applied to attach the headbar and cover the exposed skull. A layer of clear dental acrylic (Lang Dental) was then applied on top of cyanoacrylate and forms a chamber around the skull to contain the ground wire and aCSF during electrophysiological recordings. The animals received at least 3 days of rest after surgery before commencing experiments. Prior to electrophysiological recordings, we prepared a 600 μm diameter craniotomy to access the intended brain regions with Neuropixels probes.

For the groundtruth experiments (Figure 7 and 8), 100 nL of AAV2/5 CamKII-hChR2-EYFP-WPRE virus (4.6 × 10^12^ titer; UNC) was injected into ALM (2.5 mm Anterior, 1.5 mm Lateral, 0.8 mm from the dura surface) of C57BL/6J mice (n=3) (dx.doi.org/10.17504/protocols.io.bctxiwpn) in the same surgery with headbar implantation (Petreanu et al., 2009). Briefly, before the surgery, glass pipettes (Drummond Scientific Company) were pulled and sharpened to have a bevel of 35° and an opening of 20 μm at the tip. The sharpened pipette was filled from the back end with mineral oil and attached to the piston of a volumetric microinjection system (Narishige). The viral suspension was then suctioned through the tip of the pipette prior to injection. The skull over the injection site was thinned with a dental drill and punctured with the tip of the pipette. The pipette was inserted slowly (2 μm/s) to the desired depth. The virus was slowly (0.5 nL/s) injected to the desired location and the pipette was kept at the same location for 20 minutes after the end of the injection before retracted out of the brain. The virus was allowed to express for at least 4 weeks before electrophysiological recordings.

### Recordings

Electrophysiological recordings were made with Neuropixels probes (Phase 3A and B) in head-fixed mice performing an auditory delayed response task (Inagaki et al., 2018). Prior to insertion, the probe tip was painted with CM-DiI (dx.doi.org/10.17504/protocols.io.wxqffmw). Briefly, the Neuropixels probe was secured to a micromanipulator, and the back side of the probe was dipped into a 1 μL droplet of CM-DiI dissolved in ethanol (1 μg/μL). The ethanol was allowed to evaporate, and the CM-DiI was dried onto the back side of the tip. After painting with CM-DiI, the probe was attached to a micromanipulator (Sensapex) and inserted slowly (2-8 μm/s) into the brain through the craniotomy on the skull. The mean depth of recording is 3.3 mm from the surface of the brain. After reaching the desired depth for a probe, the probe was allowed to settle for 10 minutes before commencement of recording. Daily recording sessions lasted 1-2 hours, and were repeated for 3-4 days in a craniotomy. At the end of each recording session, we retracted the probe out of the brain, and cleaned the probe using Tergazyme followed by washing with distilled water. The craniotomy is sealed with removable adhesive (Kwik-Cast, World Precision Instruments) and opened again prior to the next session of recording. Within a craniotomy, we ensured each insertion is separated by at least 150 μm at the point of insertion or the insertion angles differ by more than 10°. This procedure allowed clear separation of probe tracks from different sessions of recordings within a craniotomy.

For the groundtruth experiments we stimulated ChR2-expressing neurons with a 473nm OBIS lasers (Coherent Inc.) aimed at the center of the ALM (2.5 mm Anterior, 1.5 mm Lateral) through a single mode optic fibre. The peak power was 5 mW for a 2.5 mm (4σ) spot size. We stimulated with six 2 ms square pulses spaced at 200 ms intervals, repeated every 5 seconds.

### Brain clearing

After the last recording session, mice were perfused transcardially with PBS and 4% PFA. The brains were extracted from the skull and postfixed in 4% PFA at 4°C for 12h before commencing the clearing procedure (dx.doi.org/10.17504/protocols.io.zndf5a6). Briefly, we used an alcohol-based delipidation procedure, where the brain was immersed in a 2-methyl-2-butanol (16% v/v) and 2-propanol (8% v/v) solution for 14 days. During delipidation, the brain was placed at 37°C with gentle shaking and a change of fresh solution daily. After the delipidation, the brain was washed with PBS for 1 day followed by refractive index (RI) matching in an iohexol-based solution (RI = 1.52) until it becomes visually transparent (Figure 3B) (Chi et al., 2018; Winnubst et al., 2019).

### Imaging

The cleared brain was imaged using a light-sheet microscope (Zeiss Z1). We used 5X (NA 0.1) illumination objectives and a 5X (NA 0.16) detection objective. The brain was illuminated at 488 nm (50 mW). Each horizontal section was imaged with a 150 ms exposure. Images were acquired on two imaging channels. The 504-545 nm channel captured the eYFP and autofluorescence; a channel with a 585 nm longpass filter captured CM-DiI fluorescence. Both channels were also filtered with a notch filter for the 488 nm laser emission. The 3D image of the brain (v3D) was acquired by tiling image stacks in the horizontal plane. Each image stack was 1920 μm X 1920 μm in the horizontal (XY) plane and spanned the full brain in the dorsal-ventral axis (Z). A typical brain required 20-30 stacks, with 6-12% overlap between sections in XY. The spacing between each plane within a stack was 8 μm. The size of each v3D voxel was 1.22 × 1.22 μm X 8 μm (AP x ML x DV).

After imaging the stacks were stitched using image correlation in the overlap regions (Imaris Stitcher, Bitplane). Each horizontal section was then downsampled by 5X to create the v3D used for warping to the AAT, with voxel size 6.1 × 6.1 × 8 μm.

### Template MRI brain

The MRI imaging was done using high resolution 7-T MRI at the Mouse Imaging Center at The Hospital for Sick Children in Toronto (Spencer Noakes et al., 2017). The animals were very slowly (1 mL/min) perfused with 4% PFA and MRI contrast enhancement agent Prohance (Gadoteridol, Bracco Diagnostics). After perfusion, the head was detached from the body and the skin removed from the skull. After 12h fixation with the brain inside the skull, the brains were kept in 1X PBS and 2 mM ProHance until ready to be imaged. The brains were imaged in the skull, where distortion from fixation is minimized (de Guzman et al., 2016). The resolution of the 3D stack is 40 × 40 × 40 μm.

Nine VGAT-ChR2-eYFP (JAX 014548, >P60, all male) mice contributed to the average image stack. The images of individual brains were averaged with an automated contrast based method previously described (Friedel et al., 2014; Nieman et al., 2018). Briefly, the individual images first underwent a rigid-body registration where the images are translated and rotated to be in a standard space. In the second step, individual brain images then underwent affine alignment to one target image and an averaged template was generated. The third and last step of the registration involved iterative non-linear alignment of the individual images to the averaged template in order to improve the SNR of the average (Avants et al., 2011).

Finally, the average MRI 3D was warped to the AAT using the same warping procedure as the v3D to AAT (Figure 4). A link to the MRI3D stack is available here (http://repo.mouseimaging.ca/repo/for_svoboda_hhmi/).

### Brain registration

We used a landmark-based registration package (BigWarp in ImageJ (Bogovic et al., 2016)) in which corresponding landmarks can be identified in two image stacks. The warping algorithm uses a thin plate spline for interpolation between landmarks (Duchon, 1977). Registration was done in an iterative manner. We started with a canonical set of distinct landmarks (Table 1, Supplemental Figure 4.1A) that can be reliably localized in 3D and that span the brain in all axes. After an initial round of warping, we placed additional landmarks to correct for regions with poor alignment. The warping was repeated until the 3D volume was well aligned to the template brain, judged by manual inspection. Each v3D volume required 200-300 landmarks, with emphasis on landmarks around the recording locations. Beyond the initial set of landmarks in Table 1, additional landmarks were chosen as needed for individual v3D volumes (Supplemental Figure 4.1C).

### Analyses of electrophysiological features

The extracellular voltage traces were separated into LFP and action potential (AP) bands. The LFP band signal was low-pass filtered at 300 Hz and sampled at 2.5 kHz. AP band was band-pass filtered at 300-5000 Hz and sampled at 30 kHz. For both bands of activity, before analyses, the signals from each channel was first median subtracted to remove any baseline offset from each channel. The signals across the probe then underwent common average referencing where the median across all channels on the probe at each time point was subtracted. Common average referencing is known to remove common noise across the channels on the probe (Ludwig et al., 2009).

Multi-unit activity was thresholded from the AP band at −50 μV to register events along electrode sites on the probe. We used a threshold-based algorithm, Janelia Rocket Clust (JRClust), to sort the recordings and register waveforms amongst a group of sites along the probe (Jun et al., 2017b). When possible, we also used the amplitude of the spike waveforms along the probe as electrophysiological signatures to identify transitions in brain compartments (Table 2).

For the groundtruth experiments, we fit a multi-term Gaussian model (fit function in Matlab) to the eYFP fluorescence and evoked activity along the probe (Figure 7E and 8E). The number of gaussians were specified manually based on the profile of the fluorescence and evoked activity. We compared the corresponding peak locations in fluorescence and evoked activity to quantify the accuracy of the electrode localization procedure (Figure 9).

## Terminology

Allen Mouse Common Coordinate Framework (CCF): Standard mouse brain coordinate system

Allen Anatomical Template (AAT): Image stack based on background fluorescence corresponding to the CCF (http://download.alleninstitute.org/informatics-archive/current-release/mouse_ccf/average_template/)

Allen Reference Atlas (ARA): Segmentation of the AAT into anatomical compartments (http://download.alleninstitute.org/informatics-archive/current-release/mouse_ccf/annotation/ccf_2017/)

Template MRI volume (MRI3D): MRI volume for male VGAT-ChR2-eYFP mice from the Mouse Imaging Center at The Hospital for Sick Children in Toronto (http://repo.mouseimaging.ca/repo/for_svoboda_hhmi/)

Probe: Neuropixels probe with 960 electrodes (384 recorded at the same time)

Electrode: One recording site on the Neuropixels probe

## Acknowledgements

We thank the Mouse Imaging Center at The Hospital for Sick Children in Toronto for the MRI of the brains. Andrew Recknagel, Tiago Ferreira and Jayaram Chandrashekar from the Mouselight team at Janelia for invaluable advice on brain clearing. Tim Harris Lab at Janelia for support with the Neuropixels probe recordings. The vivarium staff at the Janelia Research Campus for excellent animal care. Britton Sauerbrei for comments on the manuscript.

---

**Supplemental Figure 1.1.**
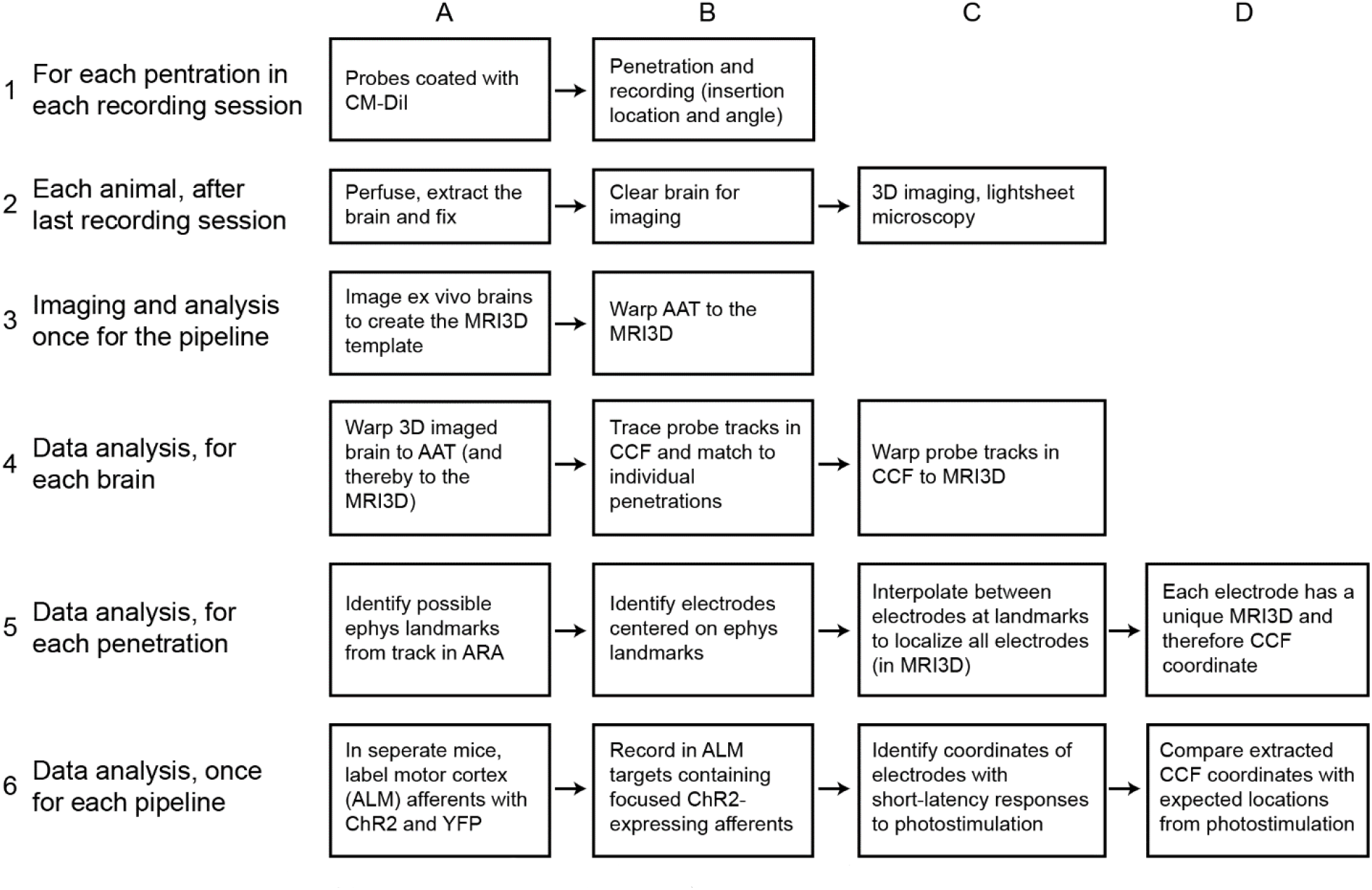
Detailed workflow for electrode localization.

**Supplemental Figure 4.1.**
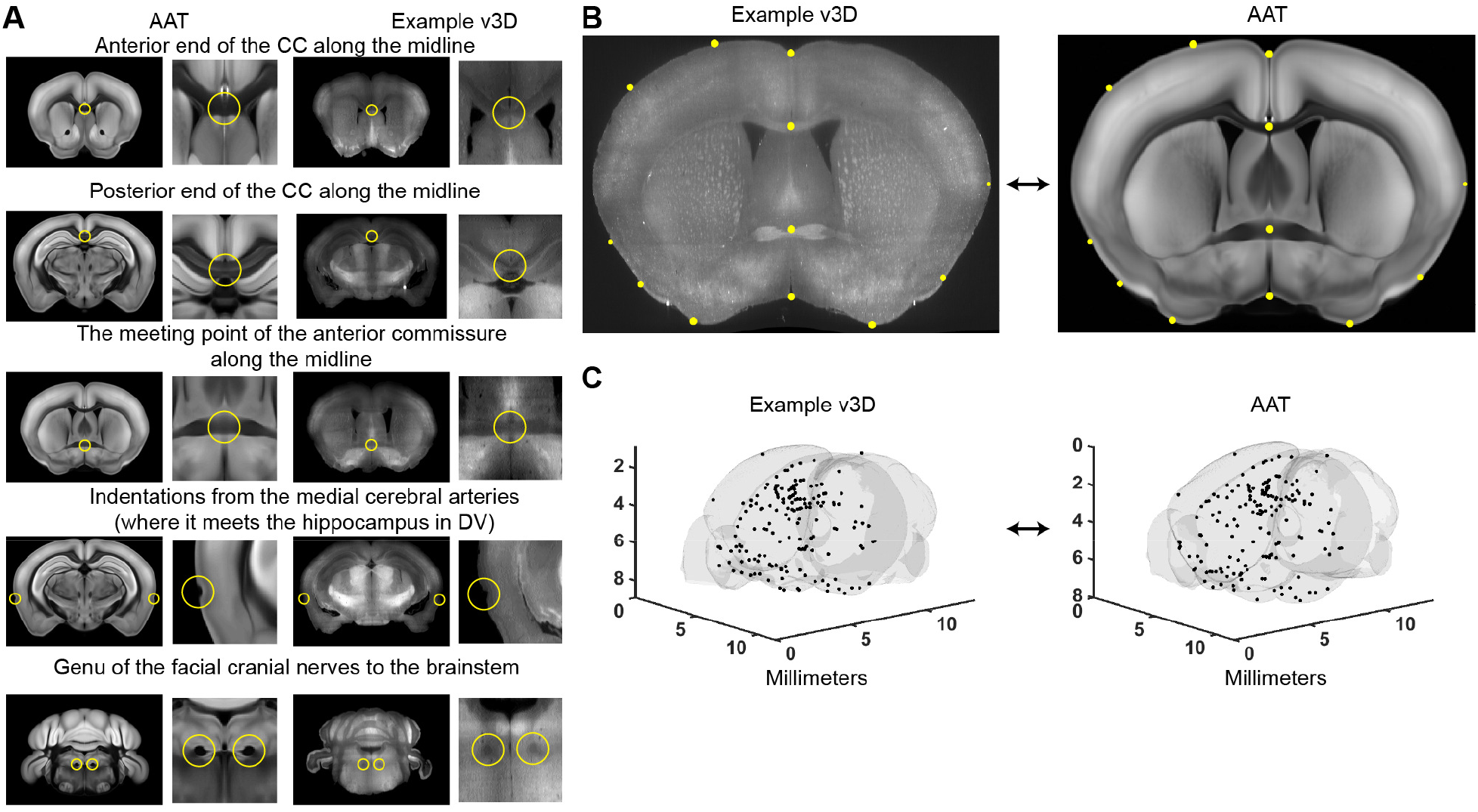
Example anatomical landmarks. (**A**) Landmarks for alignment (Table 1). The same landmarks (yellow) are identified in the AAT and v3D image volumes. (**B**) After placement of initial landmarks, warping was applied, Additional landmarks (yellow spheres) were then placed to better align the v3D and AAT. Higher densities of landmarks were placed near the probe tracks. (**C**) All landmarks of an example v3D and the AAT (black dots).

**Supplemental Figure 4.2.**
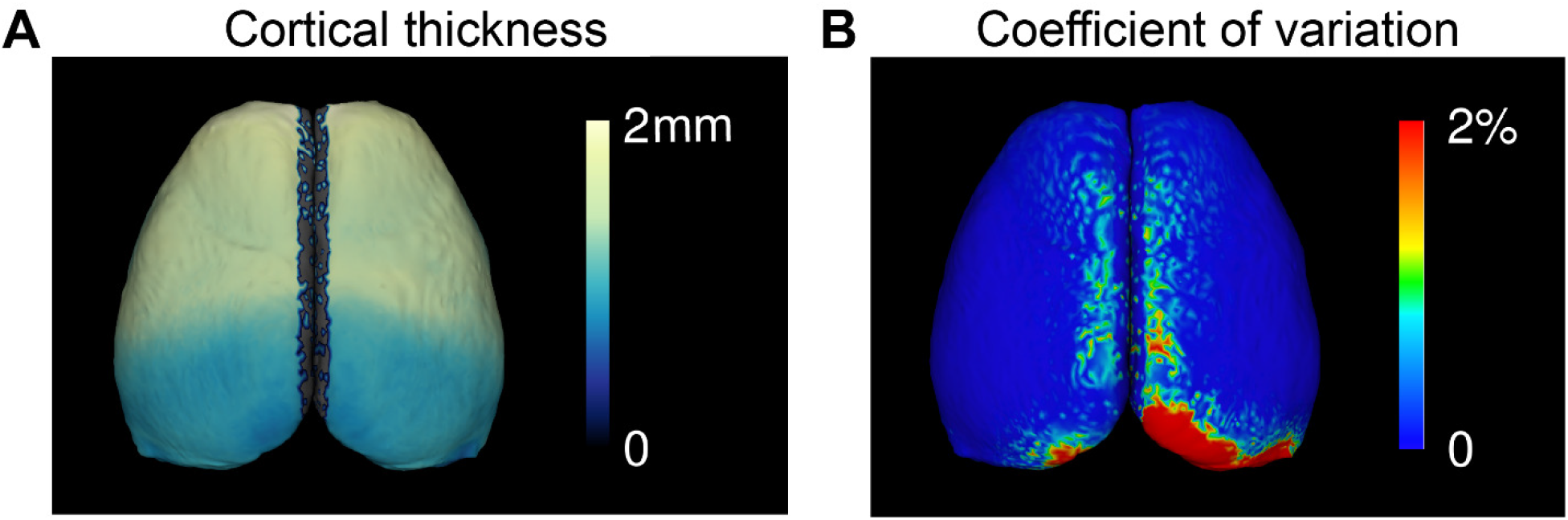
Characterization of the MRI volumes. (**A**) Mean cortical thickness measured in the nine VGAT-ChR2-eYFP mice used to generate the MRI3D template (Lerch et al., 2008). (**B**) Standard deviation of cortical thickness in MRI volumes as a percentage of the average thickness (9 mice).

**Supplemental Figure 4.3.**
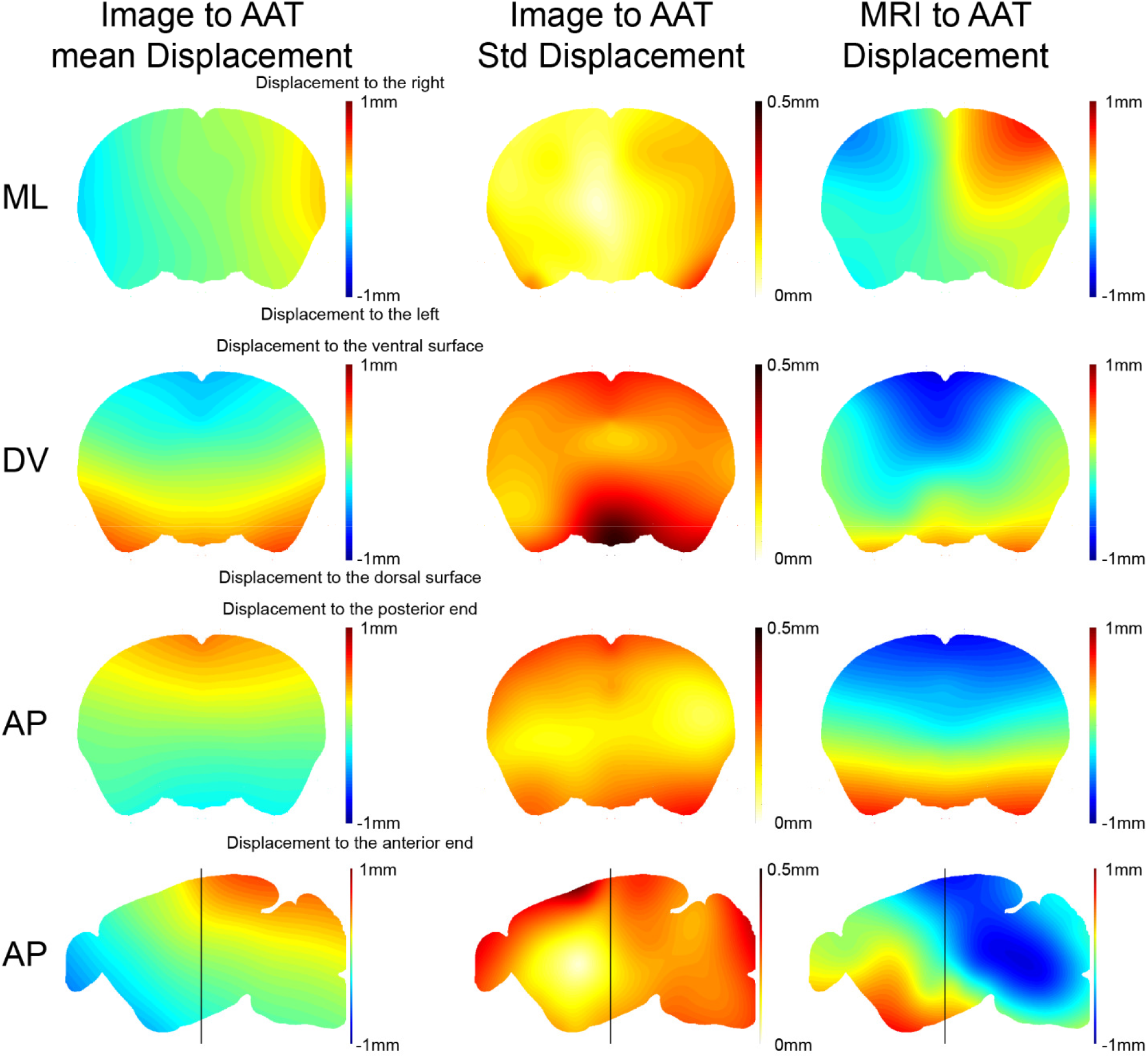
Example warp fields for one coronal section and one sagittal section. Left, averaged displacements to warp v3D image volumes onto the AAT. ML, displacements of the v3D along medial-lateral axis; DV, dorsal-ventral; AP, anterior-posterior. For example, in the top left image (ML), the v3D coronal section image has to be stretched laterally to align with the AAT (see Figure 4A). The black lines on the sagittal sections at the bottom indicate the AP position of the coronal sections. Middle, standard deviation of the displacements required to warp v3D image volumes onto the AAT (9 mice). Right, displacements to warp the average MRI3D volume onto the AAT.

